# Visualizing developmental dynamics of nuclear morphology and transport machinery in *Drosophila*

**DOI:** 10.1101/2024.10.17.618964

**Authors:** Yuki Shindo, Shruthi Balachandra, Amanda A. Amodeo

**Affiliations:** Department of Biological Sciences, Dartmouth College, Hanover, NH 03755, USA; Department of Biological Sciences, The University of Texas at Dallas, Richardson, TX 75080, USA

## Abstract

Communication between the cytoplasm and the nucleus requires a continuous exchange of molecules across the nuclear envelope (NE). The nuclear pore complex (NPC) is the gateway embedded in the NE through which cargo moves, while transport receptors mediate the passage of macromolecules through the NPC. Although their essential role as the components of the nuclear transport machinery has been extensively studied, how these factors respond to developmental and environmental cues has been underexplored. Here we tag the nucleoporin Nup96 and the transport receptor Impβ with mEGFP and mScarlet-I at their endogenous loci in *Drosophila*. We demonstrate the functionality of these markers in multiple tissues and offer new options for better visualization of nuclear morphology in densely packed, complex tissues. Then, we characterize the spatiotemporal dynamics of these markers in multiple developmental contexts. We find that Nup96 and Impβ form cytoplasmic puncta, whose size, numbers, and co-localization patterns change dynamically during oogenesis and early embryogenesis. Moreover, we find that the abundance of NPCs per nucleus decreases during early embryogenesis, complementing the emerging model in which NPCs play a regulatory role in development. The tools and observations described here will be useful in understanding the dynamic regulation of nuclear morphology and transport machinery in development.

## Introduction

The nucleus is a dynamic organelle whose composition, size, morphology, and mechanics undergo dramatic changes during development, aging, and pathogenesis (Isermann and Lammerding, 2013; Burke and Stewart, 2014; Jevtić et al., 2014; Cho and Hetzer, 2020; Hampoelz and Baumbach, 2023). The nuclear pore complex (NPC) is the gateway for macromolecular trafficking between the cytoplasm and the nucleus and is embedded in the nuclear envelope (NE) (Wente and Rout, 2010; Beck and Hurt, 2017; Lin and Hoelz, 2019). The NPC is composed of ∼30 different proteins called nucleoporins. Collectively these proteins form a selective barrier and transport channel that generates the distinct macromolecular environments of the nucleus and cytoplasm. Transport receptors, including Impβ (Importin-β), bind cargo and interact with the NPC to mediate the passage of proteins, RNAs, and other macromolecules across the NE (Chook and Süel, 2011; Wing et al., 2022). It remains poorly characterized how the nuclear transport machinery changes or maintains its homeostasis during development, when the nuclear environment is dynamically reorganized.

Visualization and segmentation of the nucleus with fluorescent protein-tagged nuclear markers are crucial for quantitative and real-time analysis of the nucleus and the molecules within (Caicedo et al., 2019). Common nuclear markers include fluorescent proteins fused to histones or nuclear localization signals (NLSs) that localize to chromatin and the nucleoplasm, respectively (Kanda et al., 1998; Shiga et al., 1996). These markers allow straightforward nuclear segmentation when cells in a tissue are positioned in 2D, such as in an epithelial monolayer. However, complex 3-dimensional tissues present additional challenges for segmentation. When cells are found in close proximity to their neighbors as in complex tissues, it is often difficult to segment individual nuclei. For example, in the *Drosophila* oocyte expressing a H2Av-mRFP histone marker, out-of-focus signal from the nucleoplasm of densely packed cells confounds attempts to segment in three dimensions (Balachandra and Amodeo, 2024). Reducing signals from the nucleoplasm while maintaining high contrast at the nuclear-cytoplasmic boundary would reduce out-of-focus background and increase the ability to segment individual nuclei. Given their enrichment at the NE and relatively low abundance in the nucleoplasm, the development of nuclear markers based on nucleoporins and transport receptors may provide an option for cleaner optical sectioning and better nuclear segmentation.

Here, we generate NE markers by endogenously tagging the nucleoporin Nup96 and the transport receptor Impβ with fluorescent proteins in *Drosophila*. We show that these markers allow for better segmentation of individual nuclei, which otherwise overlap severely when a histone marker is used. In addition, we use these reagents to visualize the developmental control of the NPC and Impβ during oogenesis and early embryogenesis. Taken together, this work provides new tools and insights into the dynamic regulation of the nuclear morphology and transport machinery in developing tissues.

## Results and Discussion

To visualize the NE in *Drosophila* tissues, we used CRISPR-Cas9 mediated genome editing to insert mEGFP (Zacharias et al., 2002) and mScarlet-I (Bindels et al., 2017) into the C-terminus of Nup98-96 (Presgraves et al., 2003) and Impβ (Ketel in *Drosophila*) (Lippai et al., 2000) at their endogenous loci, respectively (**Figs. 1A, 1B and S1**). The Nup98-96 gene is transcribed as a single transcript and its translation product is proteolytically cleaved to produce Nup98 and Nup96-mEGFP (Teixeira et al., 1997; Fontoura et al., 1999; Rosenblum and Blobel, 1999). Nup96 is a subunit of the Y-complex, one of the subcomplexes of the NPC, is present in 32 copies per NPC, and has been used as an efficient target for quantitative and super-resolution imaging of the NPC in mammalian cells (Thevathasan et al., 2019).

**Figure 1:**
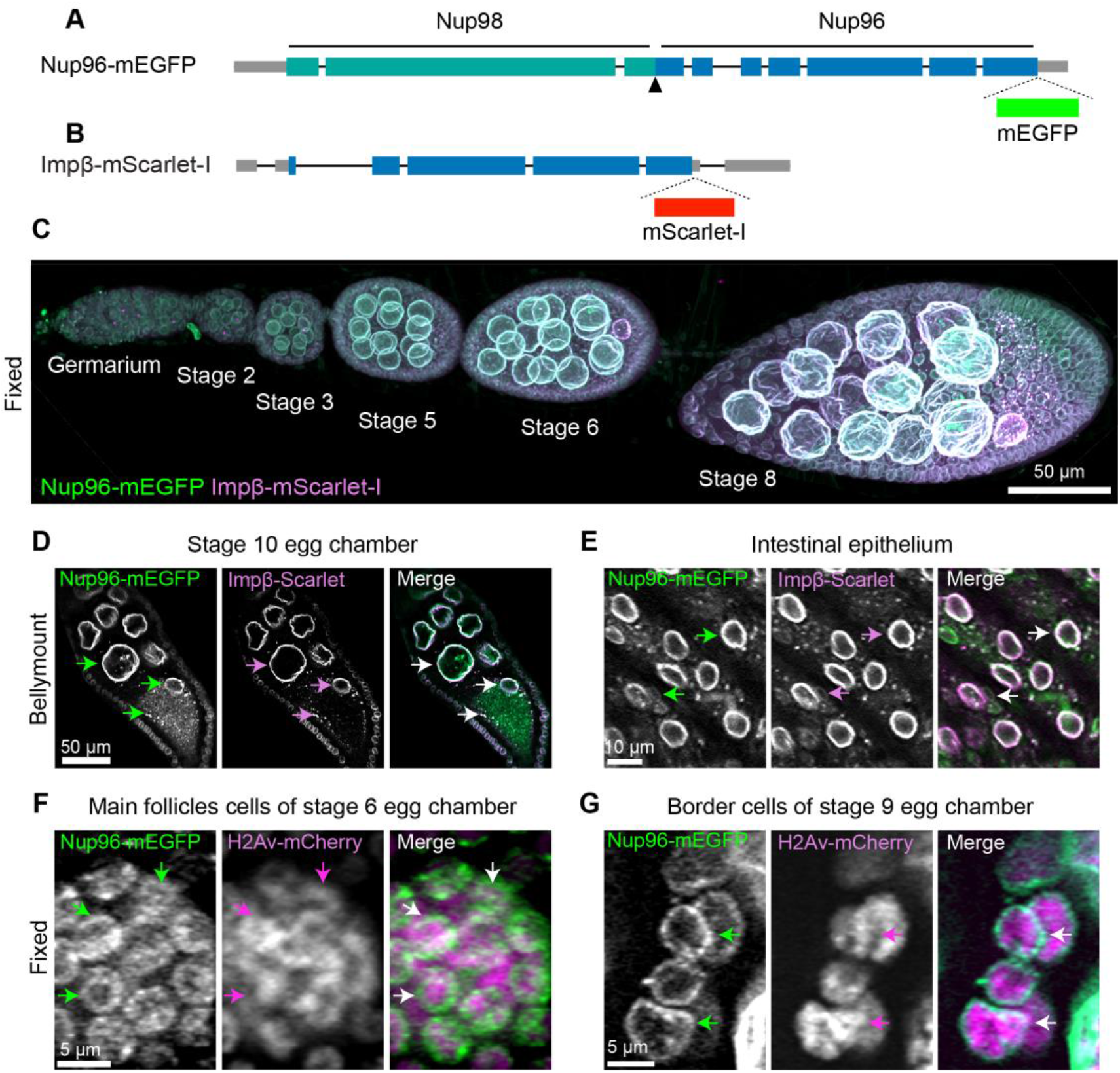
Generation of endogenously tagged Nup96-mEGFP and Impβ-mScarlet-I to visualize the nuclear morphology and transport machinery in *Drosophila*. (A) Schematic of Nup96-mEGFP at the endogenous locus. Note that the Nup98-96 gene is transcribed as a single transcript and its translation product is proteolytically cleaved to produce Nup98 and Nup96 proteins. The arrowhead indicates the cleavage site. (B) Schematic of Impβ-mScarlet-I at the endogenous locus. (C) Maximum Z-projected image of a fixed ovariole expressing Nup96-mEGFP and Impβ-mScarlet-I. Egg chambers from the germarium to stage 8 are shown. (D) Live cell imaging of stage 10 egg chamber and (E) intestinal epithelium through the abdomen of a live female using Bellymount. (F) Confocal images of fixed main follicle cells and (G) border cells expressing Nup96-mEGFP and endogenously tagged H2Av-mCherry. Individual nuclei are visible with Nup96-mEGFP even in these densely packed tissues, but not with H2Av-mCherry due to overlapping signals from neighboring cells.

Impβ is a transport receptor that, in complex with Importin-α, mediates the nuclear import of cargo with cNLSs (classical nuclear localization signals) (Adam and Geracet, 1991; Görlich et al., 1994; Wing et al., 2022). The Impα-Impβ-cargo complex binds to FG (phenylalanine-glycine) repeats of nucleoporins localized in the central channel of the NPC to pass through the NPC. Thus, Impβ is enriched at the nuclear membrane, although Impα-Impβ-cargo complexes constantly shuttle between the cytoplasm and the nucleus. In addition, it has been shown that there is a pool of Impβ that stably binds to the NPC (Lowe et al., 2015), and the stable pool likely also contributes to the enrichment of Impβ at the NE.

Both the Nup96-mEGFP and Impβ-mScarlet-I flies are homozygous viable and show no obvious defects in fertility or development. Their fluorescent signals were detected at the NE in multiple cell types tested, including embryos, ovarian germline and somatic cells, and intestinal epithelial cell types such as enterocytes and progenitors in both fixed tissues and intact animals (**Figs. 1C–1G and 2**), suggesting that they can be used as ubiquitously expressed nuclear markers. We note that these markers do not lead to overexpression of the tagged proteins unlike previously reported NE markers and thus provide a more physiological context. This is critical since changes in the abundance of nuclear transport machinery can dramatically affect nuclear and developmental parameters (Levy and Heald, 2010; Jevtić and Levy, 2015).

We sought to test if the NE markers we generated could improve the visualization of nuclei in densely packed tissues. To this end, we focused on the egg chamber in which multiple cell types of different sizes are closely opposed and overlap in three dimensions. Follicle cells form an epithelium that encloses the oocyte and 15 nurse cells and ultimately generate the egg shell during oogenesis (Bastock and Johnston, 2008). Nuclear segmentation of individual follicle cells is challenging due to the overlap of fluorescence signal from surrounding nuclei when imaged with a histone H2Av marker (**Fig. 1F and Fig. S2**). Our NE markers allowed segmentation of individual follicle cell nuclei in conditions where we could not segment the same nuclei with a histone marker (**Fig. 1F**). To further test the segmentation improvements generated by our new markers, we visualized the nuclei from the cluster of 6-10 collectively migrating border cells (**Fig. 1G**). Segmenting border cells in 3D is often challenging and frequently involves multiple markers for the cell membranes and nucleus (Cliffe et al., 2017). We found a similar improvement in visualizing individual nuclei using our NE markers (**Fig. 1G**). Therefore, our NE markers can improve the nuclear segmentation in complex 3D tissues.

Next, we sought to utilize these markers to study the regulation of the nuclear transport machinery. We visualized the dynamics of Nup96-mEGFP throughout the early syncytial mitotic cell cycle in early *Drosophila* embryos. We observed that Nup96-mEGFP signal decreased at the NE at the onset of mitosis but remained associated with the spindle envelope structure as nuclei underwent semi-closed mitosis (**Fig. 2A**). Overall, Nup96-mEGFP showed similar dynamics to that of the Nup107 transgenes (Katsani et al., 2008; Ren et al., 2019), consistent with the fact that both Nup96 and Nup107 are subunits of the Y-complex.

**Figure 2:**
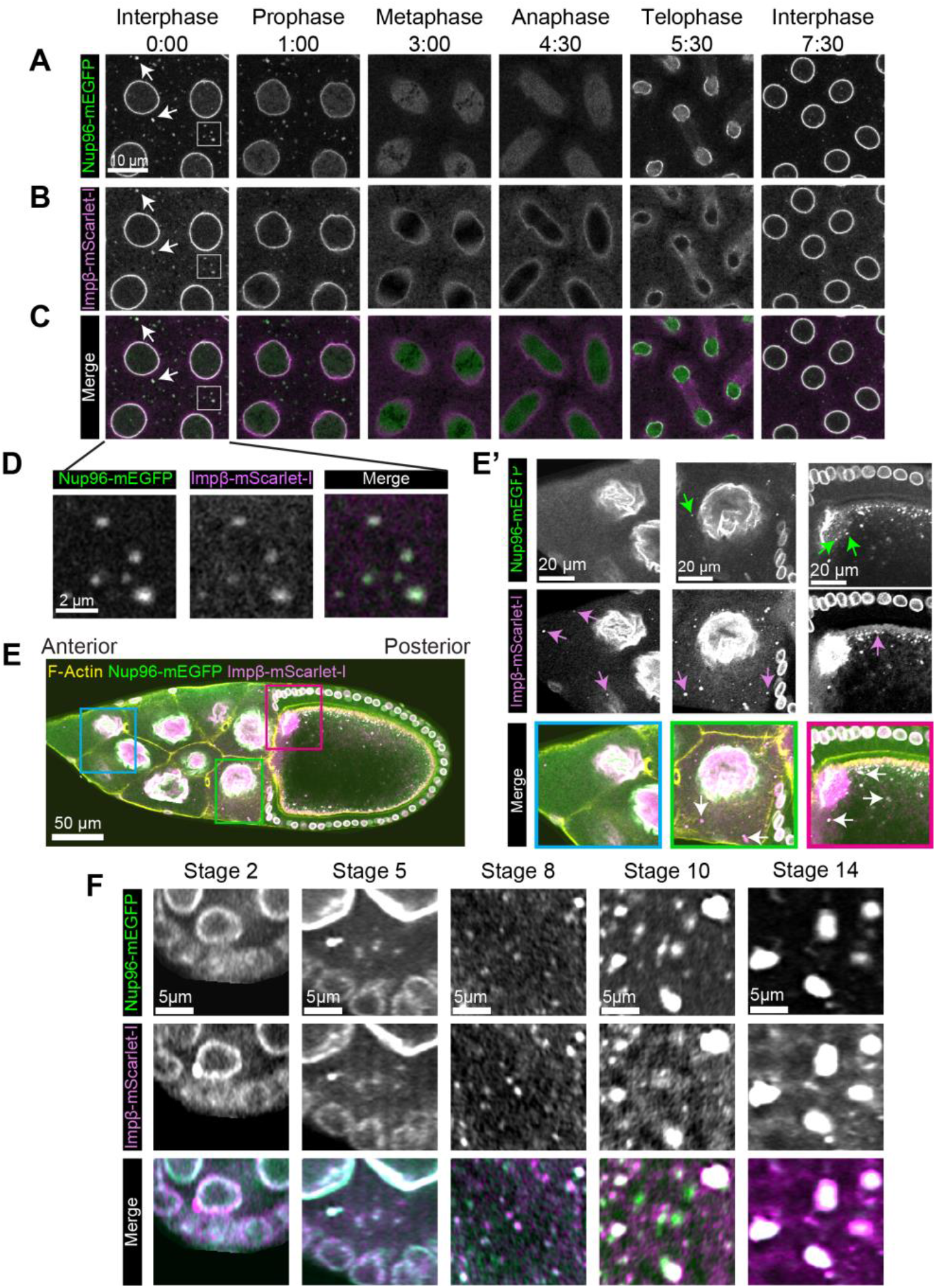
Temporal dynamics of Nup96 and Impβ during the mitotic cell cycle and oocyte development. (A)–(C) Timelapse confocal images of Nup96-mEGFP and Impβ-mScarlet-I in cycle 11 in early embryos. Arrowheads indicate cytoplasmic puncta of Nup96-mEGFP and Impβ-mScarlet-I. (D) Enlarged images of panels A–C (gray box), showing colocalization of Nup96-mEGFP and Impβ-mScarlet-I puncta. (E) Confocal image of a fixed stage 10 egg chamber expressing Nup96-mEGFP and Impβ-mScarlet-I, stained for F-actin. Enlarged images of the regions labeled with the magenta (anterior nurse cells), blue (posterior nurse cells), and green (oocyte) boxes are shown in (E’). Impβ-mScarlet-I puncta are observed in the oocyte and all nurse cells across the anterior-posterior axis, whereas Nup96-mEGFP puncta are localized only in the oocyte and nurse cells in direct contact with the oocyte. (F) Temporal dynamics of Nup96-mEGFP and Impβ-mScarlet-I puncta in the oocyte cytoplasm during oocyte development. The size and number of these cytoplasmic puncta in the oocyte change over developmental time.

In addition to the NE signals, we observed Nup96-mEGFP puncta in the cytoplasm (**Fig. 2A, arrowhead**). These puncta likely represent nucleoporins stored in a subdomain of the ER called the annulate lamellae (AL) (**Figs. S3A–3D**) (Stafstrom and Staehelin, 1984; Eymieux et al., 2021). The Nup96-mEGFP puncta persisted during the interphase and prophase, but almost completely dissolved in metaphase, followed by reassembly in telophase or early interphase (**Fig. 2A**). For comparison, we also visualized emGFP-Nup358, which has been shown to form cytoplasmic puncta in early *Drosophila* embryos (Hampoelz et al., 2019, 2016). We observed that emGFP-Nup358 puncta decreased in size during mitosis but did not disappear completely (**Fig. S3E**). The distinct dynamics of Nup96 and Nup358 puncta suggest a hierarchical nature of nucleoporin puncta organization, consistent with the previous observation that Nup358 acts as the seed of the punctate structure on which other components are decorated.

Interestingly, we found that Impβ-mScarlet-I also forms cytoplasmic puncta in early embryos (**Fig. 2B**). Similar to Nup96-mEGFP puncta, the Impβ-mScarlet-I puncta persisted throughout interphase and prophase, disappeared during metaphase and anaphase, and reformed at the end of mitosis. The Impβ-mScarlet-I puncta overlapped with Nup96-mEGFP puncta in the cytoplasm (**Figs. 2C and 2D**), suggesting that AL are composed not only of nucleoporins but also of transport receptors (Raghunayakula et al., 2015). To test the presence of Impβ puncta in other tissues and to gain insight into their biogenesis and NE assembly, we turned to the ovary in which nucleoporin puncta have been shown to form (Hampoelz et al., 2019). Indeed, we observed accumulation of both Nup96-mEGFP and Impβ-mScarlet-I puncta in the oocyte cytoplasm during oogenesis (**Fig. 2E**). Impβ-mScarlet-I puncta were also found in the cytoplasm of most nurse cells. However, Nup96-mEGFP puncta were restricted to the most posterior nurse cells. Where they co-occurred, Nup96-mEGFP and Impβ-mScarlet-I were mostly colocalized (**Fig. 2E**, stage 10, **Fig. S4**). These distinct and overlapping behaviors of Nup96-mEGFP and Impβ-mScarlet-I puncta across the spatial coordinate of the egg chamber suggest that there are multiple and/or stepwise pathways for the formation of the puncta.

In addition to the spatial patterning of Nup96 and Impβ puncta along the anterior-posterior axis of the egg chamber, we found changes in the number and size of the cytoplasmic puncta over developmental stages during oogenesis. Both Nup96-mEGFP and Impβ-mScarlet-I puncta started to accumulate in the oocyte cytoplasm as early as stage 5 of oogenesis (**Fig. 2F**), similar to Nup107 as described previously (Hampoelz et al., 2019). The number of the puncta in the oocyte cytoplasm increased with the progression of oogenesis until stage 8, after which the puncta increased in size rather than number towards the end of oogenesis (**Fig. 2F**). Overall, both Nup96-mEGFP and Impβ-mScarlet-I puncta in the oocyte cytoplasm behaved similarly during oogenesis. However, there was a notable difference in the behavior of Nup96 and Impβ at the nurse cell NE, where Impβ-mScarlet-I levels decreased after nurse cell dumping while Nup96-mEGFP levels remained unchanged (**Fig. S4**). The stable association of Nup96 at the nurse cell NE may be important for the proper completion of oogenesis, as Nup154, another scaffold nucleoporin, is known to play an important role in nurse cell elimination following dumping (Riparbelli et al., 2007).

Our observations that the behavior of Nup96 and Impβ puncta changes dynamically during oogenesis prompted us to visualize their dynamics in other developmental contexts. We were particularly interested in early embryogenesis because we have previously proposed a possible role for the nuclear transport machinery in early embryonic cell cycles (Shindo and Amodeo, 2021). The large stockpile of nucleoporins and transport receptors produced during oogenesis would be important for early embryogenesis, during which the demand for nuclear components increases exponentially as the embryo undergoes the reductive cleavage division cycles. We focused on the syncytial blastoderm stage (cycles 10-13), during which there is little zygotic transcription and nuclei divide rapidly without intervening cytokinesis (Blythe and Wieschaus, 2015; Harrison and Eisen, 2015). Using time-lapse imaging, we observed Nup96-mEGFP puncta over multiple cycles and found that the number and size of the puncta decreased dramatically over the course of the syncytial blastoderm stage (**Fig. 3A**). At cycle 13 interphase, there were almost no detectable Nup96-mEGFP puncta in the cytoplasm. The decrease in the abundance of cytoplasmic Nup96-mEGFP puncta is likely due, at least in part, to the increased demand for NE components during the syncytial blastoderm stage where repeated cleavage divisions rapidly dilute pre-existing NPCs at the NE into daughter nuclei.

**Figure 3:**
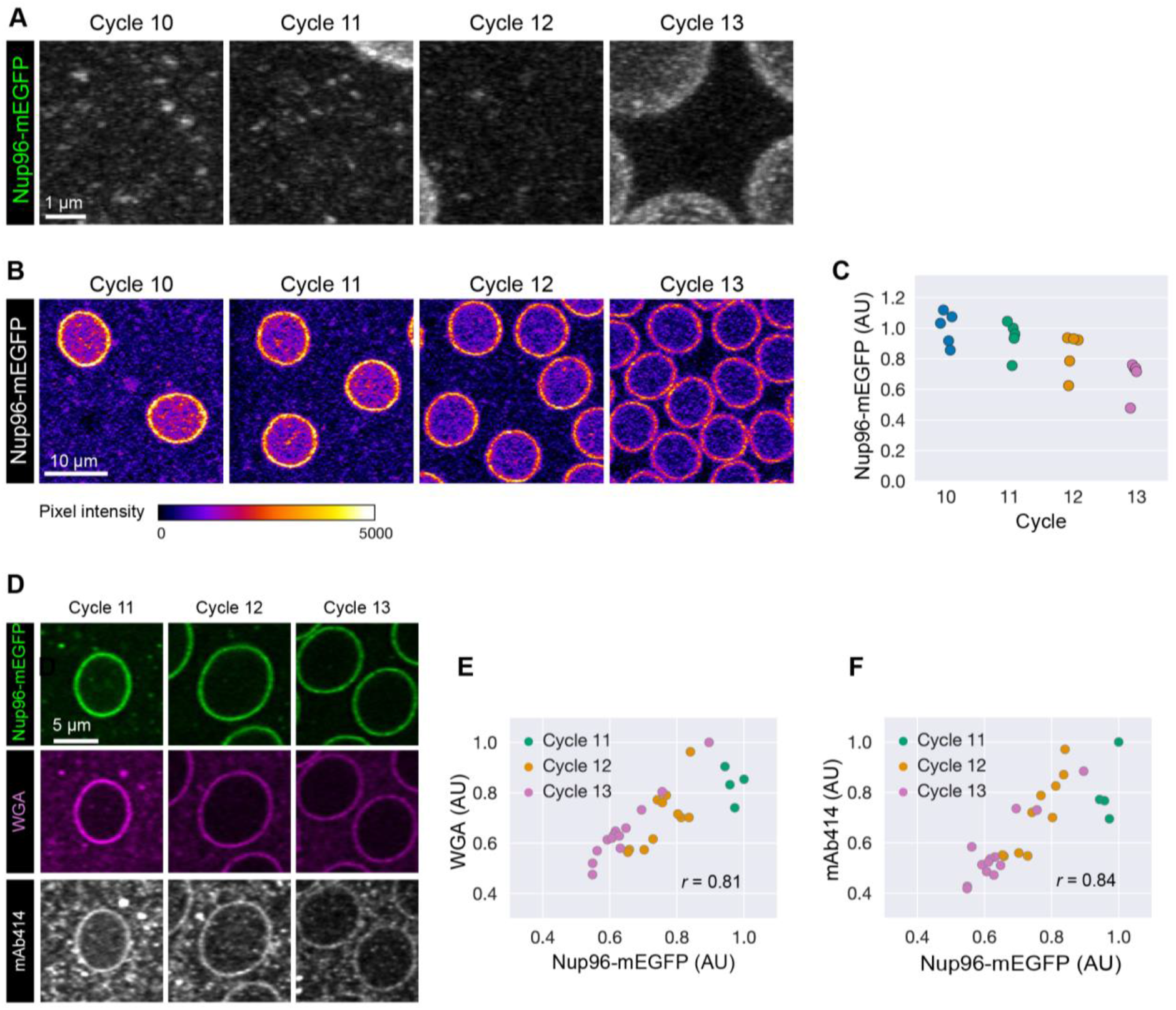
Dynamic reduction in Nup96-mEGFP levels during early embryogenesis. (A) Time-lapse confocal images of Nup96-mEGFP cytoplasmic puncta during the syncytial blastoderm stage. (B) Time-lapse confocal images of Nup96-mEGFP at the NE over the course of the syncytial blastoderm stage. (C) Average fluorescence intensities of Nup96-mEGFP at the NE, showing a reduction in Nup96-mEGFP levels at the NE during early embryogenesis. n = 5 embryos. (D) Confocal images of nuclei in fixed embryos with WGA and mAb414 staining. (E) and (F) Average fluorescence intensities of Nup96-mEGFP, WGA, and mAb414 at the NE. Their intensities decrease with embryonic developmental progression from cycles 11 to 13. n = 28 embryos.

Finally, we asked how the levels of Nup96-mEGFP at the NE correlate with the dynamics of the cytoplasmic puncta. Using time-lapse imaging, we found that per-nucleus levels of Nup96-mEGFP at the NE indeed decreased from cycle 10 to cycle 13 (**Figs. 3B and 3C**). Our observation suggests that while the cytoplasmic pool may help to counteract the loss of Nup96 at the NE with each division cycle, it becomes depleted with developmental progression. That is, the balance between rapid dilution due to the cleavage divisions and new supply from the cytoplasmic pool of nucleoporins dynamically sets the abundance of NPCs at the NE during early embryogenesis. Because Nup96 is a scaffold nucleoporin critical for NPC assembly (Walther et al., 2003), the reduction in Nup96 levels at the NE suggests the decrease in the density and number of NPCs. To test this possibility, we stained fixed embryos with CF568-conjugated wheat germ agglutinin (WGA) and the mAb414 antibody that binds several FG-nucleoporins. We observed that WGA and mAb414 signals at the NE also decrease during cycles 11-13 (**Figs. 3D–F**), consistent with the idea that the per-nuclear abundance of NPCs, not specifically Nup96, decreases during early embryonic development.

While the NPC has been thought to be relatively invariant across cell types because of its essential housekeeping role, our data show that the number of NPCs changes dynamically during early embryonic development in *Drosophila*. This observation complements the recent finding in zebrafish that the nucleoporin composition of NPCs changes during early embryogenesis (Shen et al., 2022). Associations between the number of NPCs and cell state have also recently been observed in other cellular contexts, including heart development and cancer survival (Han et al., 2022; Sakuma et al., 2021). Together, these findings suggest a role for the nuclear transport machinery as an active regulator of cell physiology and development. It remains to be determined how these changes in the number and composition of NPCs affect cellular processes and development. In zebrafish, it has been proposed that the changes in the NPC composition control the nuclear entry of transcription factors important for zygotic genome activation (Shen et al., 2022). Whether changes in NPC state specifically affect a subset of cargoes or if they have a global effect on cargo nuclear transport remains unclear. In addition, NPCs are also known to interact with chromatin to regulate gene expression and chromatin organization (Pascual-Garcia and Capelson, 2019; Cho and Hetzer, 2020). Understanding how these changes in NPCs affect such off-pore roles requires further study.

In conclusion, we have presented endogenously-tagged NE markers that can allow better segmentation of individual nuclei in densely packed tissues in *Drosophila*. We have used this tool to visualize the dynamics of Nup96 and Impβ in the ovary and embryo, finding the spatiotemporal organization of Nup96 and Impβ puncta in the cytoplasm during oogenesis and the dynamic reduction in the number of NPCs during early embryogenesis. Taken together, our tools and observations presented in this work have set the stage for further interrogation of the nuclear dynamics, biosynthesis of nuclear transport machinery, and their roles in development.

## Materials and methods

### Contact for reagent and resource sharing

Further information and requests for resources and reagents should be directed to Yuki Shindo.

### Fly husbandry

All fly stocks were maintained at room temperature of approximately 22 °C on standard molasses media. Embryos were collected on yeasted apple juice agar plates for two hours at 25°C and dechorionated with 4% sodium hypochlorite for 2 min followed by a wash in deionized water.

### Plasmids and transgenesis

The pU6-BbsI-chiRNA and pScarless-sfGFP-3xP3-DsRed plasmids were a gift from Melissa Harrison & Kate O’Connor-Giles & Jill Wildonger (Addgene plasmid #45946 and #80811). To generate endogenously tagged Nup96-mEGFP, Impβ-mScarlet-I, and H2Av-mCherry flies, single CRISPR target sites near the stop codon of the endogenous gene loci were selected using Target Finder (Gratz et al., 2014) and the designed target gRNA sequences (Nup96: TGCTGCCGCTCGCTAAATGG; Impβ: TACCCAGGTCATCACGCAGT; H2Av: GGTGCAGGATCCGCAGCGGA) were subcloned into the pU6-BbsI-chiRNA vector. For Nup96-mEGFP, a 928 bp left homology arm with a synonymous mutation at the PAM site, followed by a linker (TSRRYRGPGIHRPVAT) and mEGFP (Thevathasan et al., 2019), and a 942 bp right homology arm were synthesized and assembled in the pScarlessHD plasmid backbone. For Impβ-mScarlet-I, sfGFP in the pScarless-sfGFP-3xP3-DsRed plasmid was replaced with mScarlet-I and then a 1 kb left homology arm and a 1 kb right homology arm were synthesized and assembled in the pScarless-mScarlet-I-3xP3-DsRed plasmid backbone. For H2Av-mCherry, a 1 kb left homology arm with a synonymous mutation at the PAM site and a 1 kb right homology arm were synthesized and assembled in the pScarless-Dendra2 plasmid backbone (Shindo and Amodeo, 2019), and then Dendra2 was replaced by mCherry. The plasmids of gRNA and homology arms were co-injected into nos-Cas9 embryos (TH00787.N for Impβ-mScarlet-I and TH00788.N for Nup96-mEGFP and H2Av-mCherry) (BestGene and Rainbow Transgenic Flies) and DsRed+ progeny were screened. The DsRed marker was removed through a cross to nos-PBac flies (a gift from Robert Marmion and Stanislav Shvartsman) (Keenan et al., 2020). Insertion of the fluorescent proteins was confirmed by PCR and Sanger sequencing. Expression of the tagged proteins was confirmed by Western blotting with anti-GFP or anti-RFP antibodies (**Fig. S1**).

### Ovary fixation and staining

Ovaries from 8 to 15 females were dissected in PBSTw (PBS with 0.1% Tween-20) and fixed in a 1:1 mixture of 4% formaldehyde (Thermo Scientific, 28908) in PBS and heptane (Sigma Aldrich, 34873) for 20 minutes at room temperature. Following fixation, samples were washed three times for 10 minutes each in PBSTw. Subsequently, samples were permeabilized in 1% Triton-X100 (Sigma Aldrich, 9036-19-5) for 1 hour and blocked in 1% BSA (Millipore-Sigma, A9647) prepared in PBSTw for an additional hour. Samples were then stained with a mixture of Hoechst (1:1000) (Thermo Scientific, 33342) and Phalloidin (1:500) (Thermo Fischer Scientific, A22283) prepared in the blocking solution for 3 hours at room temperature on a nutator. After staining, samples were given three quick washes and two 15-minute washes in PBSTw. Finally, samples were mounted in EverBrite mounting media (Biotium, #23001) with a 0.3 mm SecureSeal™ Imaging Spacer (Grace Bio Labs, SS1X20) and sealed with transparent nail polish.

### Live imaging of egg chambers and gut (Bellymount)

Egg chambers and gut tissues in live females were observed using Bellymount (Koyama et al., 2020; Balachandra and Amodeo, 2024). Briefly, four to five-day-old healthy females were anesthetized on a fly CO2 pad for mounting. Five females were affixed to the glass surface of a 50 mm MatTek dish (MatTek, P50G-1.5-30-F) using clear Elmer’s Liquid School Glue (Elmer’s, E309). An anesthetized female was carefully positioned on the glue and gently pressed with blunt forceps. The fly was secured by placing a 0.5 cm^2^ compression glass piece starting from her thorax (cut from a 24 × 60 mm cover glass; VWR International, 48393-106). A pair of 2 mm^2^ double-sided adhesive spacers, 0.36 mm thick (Millipore-Sigma, GBL620003-1EA), were positioned on either side of the fly between the compression glass and the dish surface. A small amount of yeast paste was provided near the fly’s legs to stimulate feeding. Flies were anesthetized with CO2 while imaging to stabilize the fly movement.

### Embryo fixation and staining

Embryos were collected for 2.5 h, developed for 30 min, dechorionated, and fixed in a 1:1 mixture of 4% formaldehyde and heptane for 20 min and devitellinized in a 1:1 mixture of methanol and heptane followed by 3×washes in methanol. The fixed embryos were rehydrated by 3× washes in PBSTw (PBS with 0.05% Tween-20) and blocked with Image-iT FX Signal Enhancer (Thermo Fisher, I36933) for 30 min at room temperature. Blocked embryos were incubated with the mouse anti-Nuclear Pore Complex Proteins (1:200, BioLegend, #902901) antibody in PBSTw supplemented with 5% NGS (Normal Goat Serum, Cell Signaling Technology, #5425) overnight at 4 °C. Embryos were washed in PBSTw twice for 10 min each and then incubated with Alexa Fluor 647 F(ab’)2-Goat anti-Mouse IgG (1:300, Invitrogen, A-21237) in PBSTw 5% NGS for 1 h at room temperature in the dark. At 40 min into the secondary antibody incubation, Hoescht 33342 (1:5000, Thermo Fischer Scientific, #62249) and WGA (Wheat Germ Agglutinin)-CF568 (1:300, Biotium, 29077) were added. Embryos were washed in PBSTw four times for 10 min each and mounted on glass slides with SecureSeal™ Imaging Spacer in EverBrite Mounting Medium.

### Microscopy

All the imaging experiments presented in this paper were carried out with a Zeiss LSM980 Airyscan 2 microscope at room temperature of approximately 22 °C. Raw images were Airyscan-processed using ZEN 3.3. The following lasers were used: 405 nm for Hoescht 33342; 488 nm for mEGFP or emGFP; 561 nm for mScarlet-I, mCherry, mRFP, or CF568; and 647 nm for Alexa Fluor 647.

Egg chambers and gut tissues in either fixed samples or live females were imaged with a 20×0.80 NA air objective. Z-stack images (0.124-0.149 µm/pixel, approximately 80 µm depth, 0.5-1 µm Z-step size) were acquired with the Multiplex CO-8Y mode.

For live imaging of embryos, embryos were collected for 2 h, dechorionated, mounted in deionized water on a glass-bottom microwell dish (MatTek), and imaged with a 40× 1.30 NA oil immersion objective. For imaging of Nup96-mEGFP, RFP-KDEL, emGFP-Nup358, and Impβ-mScarlet-I during the mitotic cell cycle, time-lapse images (0.046 µm/pixel) were acquired with the Multiplex SR-4Y mode at a time resolution of 6 s with or without Z-stacks (7 Z-slices, 1 µm Z-step size). For imaging of Nup96-mEGFP throughout the syncytial blastoderm stage, time-lapse Z-stack images (0.092 µm/pixel, 16 Z-slices, 1 µm Z-step size) were acquired with the Multiplex CO-8Y mode at a time resolution of 1 min. The NE was segmented using Labkit (Arzt et al., 2022) implemented in Fiji/ImageJ, and fluorescence intensities at the NE were quantified using Fiji/ImageJ.

For imaging of fixed embryo samples, fluorescence images (0.076 µm/pixel) were acquired with a 40× 1.30 NA oil immersion objective. The Multiplex CO-8Y mode was used. Z-slices were collected at a step size of 0.5 µm. Image analysis including the segmentation of the NE and measurements of fluorescence intensities at the NE was performed using custom Python scripts.

### Western blotting

Either embryos collected over a one-hour period or ovaries dissected from females fed with yeast paste were lysed in ice-cold RIPA buffer (Sigma, R0278) supplemented with 1x protease inhibitor cocktail (Sigma, P2714). Proteins were separated on a 7.5% TGX Stain-Free polyacrylamide gel (Bio-Rad), crosslinked for 45 s under UV, and transferred to a low fluorescence PVDF membrane. Membranes were blocked with 5% low-fat milk in PBST, incubated overnight in primary antibodies at 4 °C, washed, and incubated in secondary antibodies for 1 hour at room temperature. Chicken anti-GFP (Aves Labs, GFP-1010; 1:1000), mouse anti-RFP (proteintech, 6g6; 1:2000), goat anti-chicken IgY CF647 (Biotium, #20044; 1:1000), and goat anti-mouse IgG Alexa Fluor Plus 647 (Invitrogen, A32728; 1:2000) were used. Fluorescence was detected using a gel imager (Bio-Rad ChemiDoc MP).

### Fly stocks

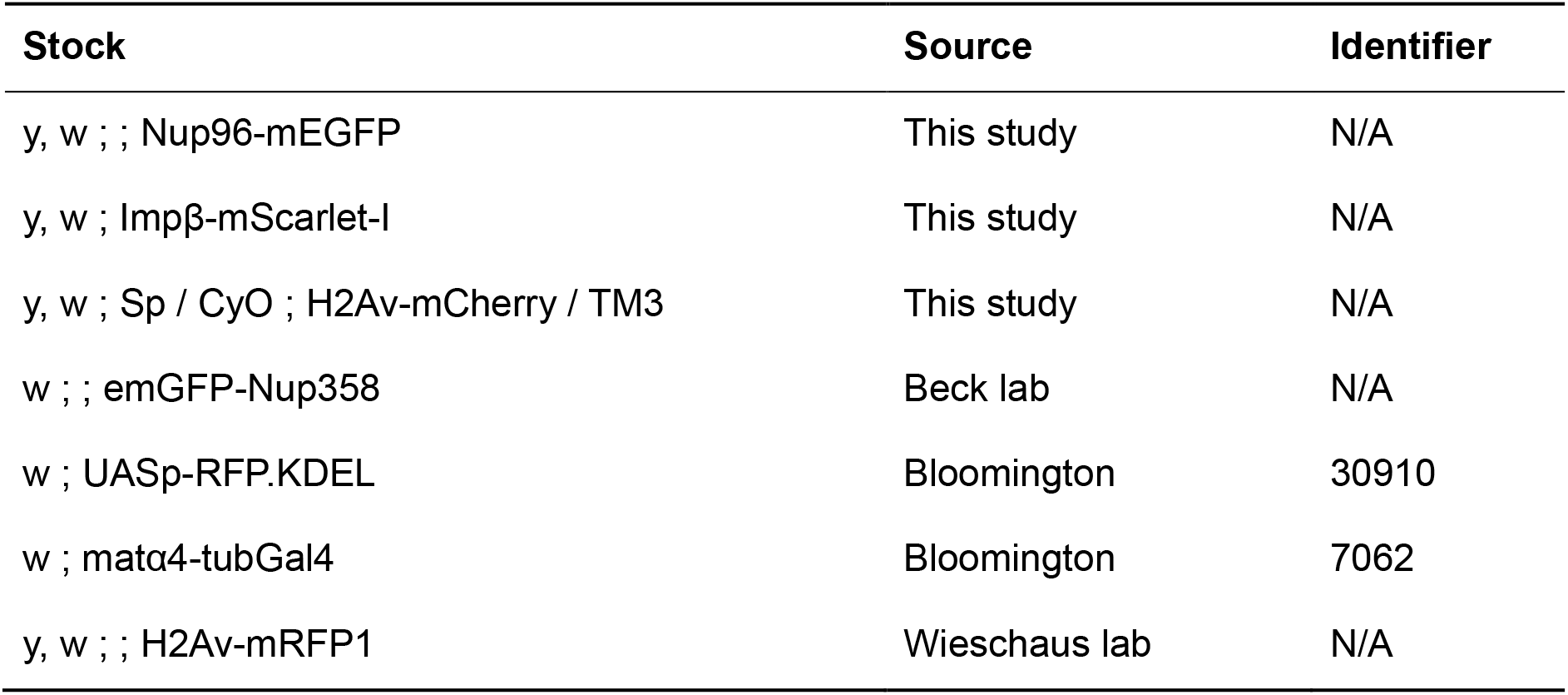

## Acknowledgments

We thank Melissa Harrison, Kate O’Connor-Giles, and Jill Wildonger for plasmids; Robert Marmion, Stanislav Shvartsman, Martin Beck, and the Bloomington Drosophila Stock Center (National Institutes of Health P40OD018537) for fly stocks. We also thank Ann Lavanway, Dartmouth bioMT, Britton Johnson, and *Drosophila* Media Core Facility at Dartmouth College. Y.S. was supported by the Uehara Memorial Foundation and the Osamu Hayaishi Memorial Scholarship for Study Abroad. This work was supported by NIH NIGMS MIRA (1R35GM150853-01) and NIH NIGMS COBRE award (P20-GM113132).

## Author Contribution

Conceptualization, Y.S. and S.B.; Investigation, Y.S. and S.B.; Writing – Original Draft, Y.S. and S.B.; Writing – Review and Editing, Y.S., S.B., and A.A.A.

## Supplemental Figures

**Figure S1:**
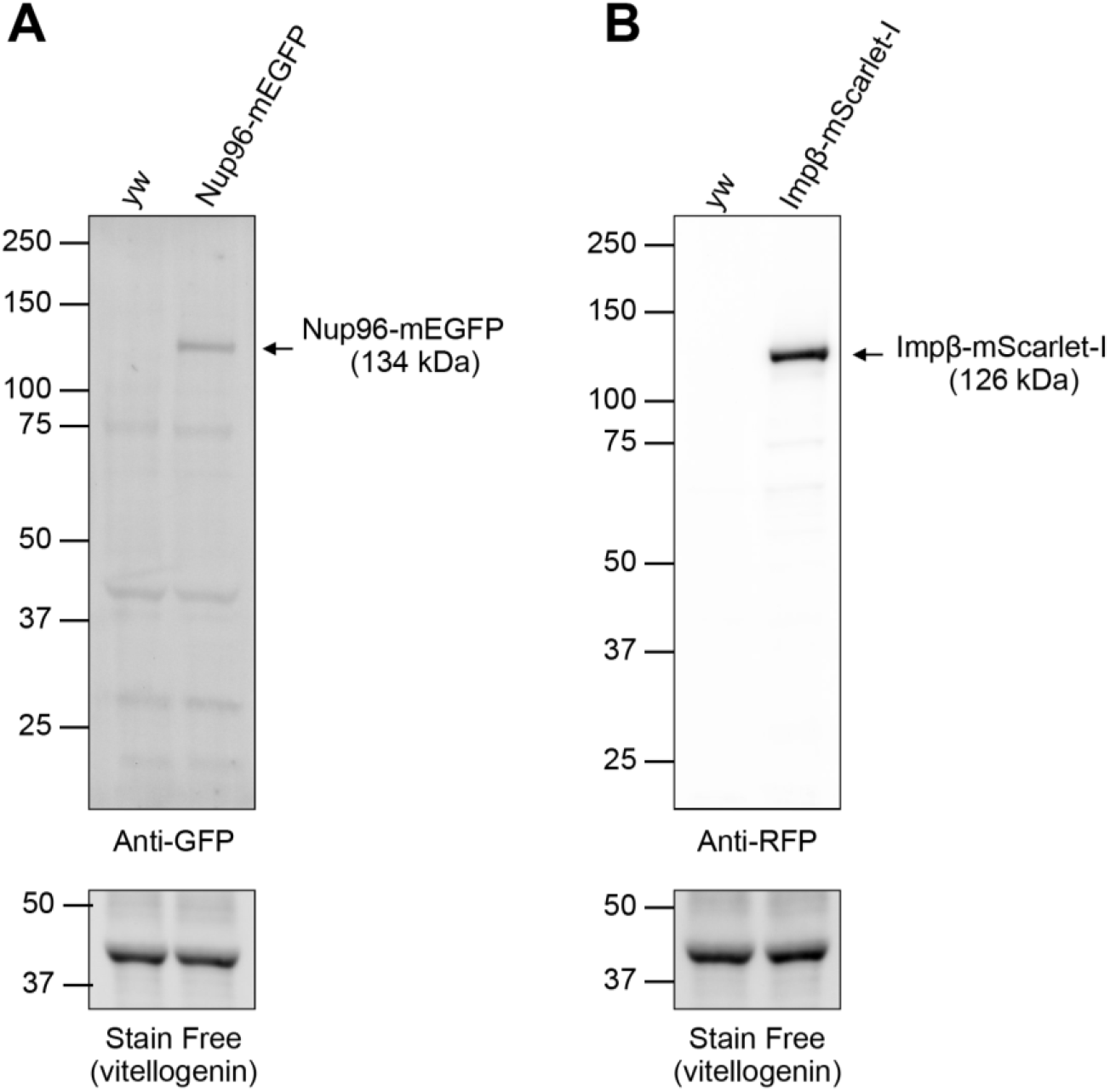
Validation of endogenously-tagged Nup96-mEGFP and Impβ-mScarlet-I. Embryos were collected over a one-hour period and total embryo lysates were analyzed by western blotting. The specific bands consistent with the predicted sizes of (A) Nup96-mEGFP (134 kDa) and (B) Impβ-mScarlet-I (126 kDa) were detected with anti-GFP and anti-RFP antibodies, respectively.

**Figure S2:**
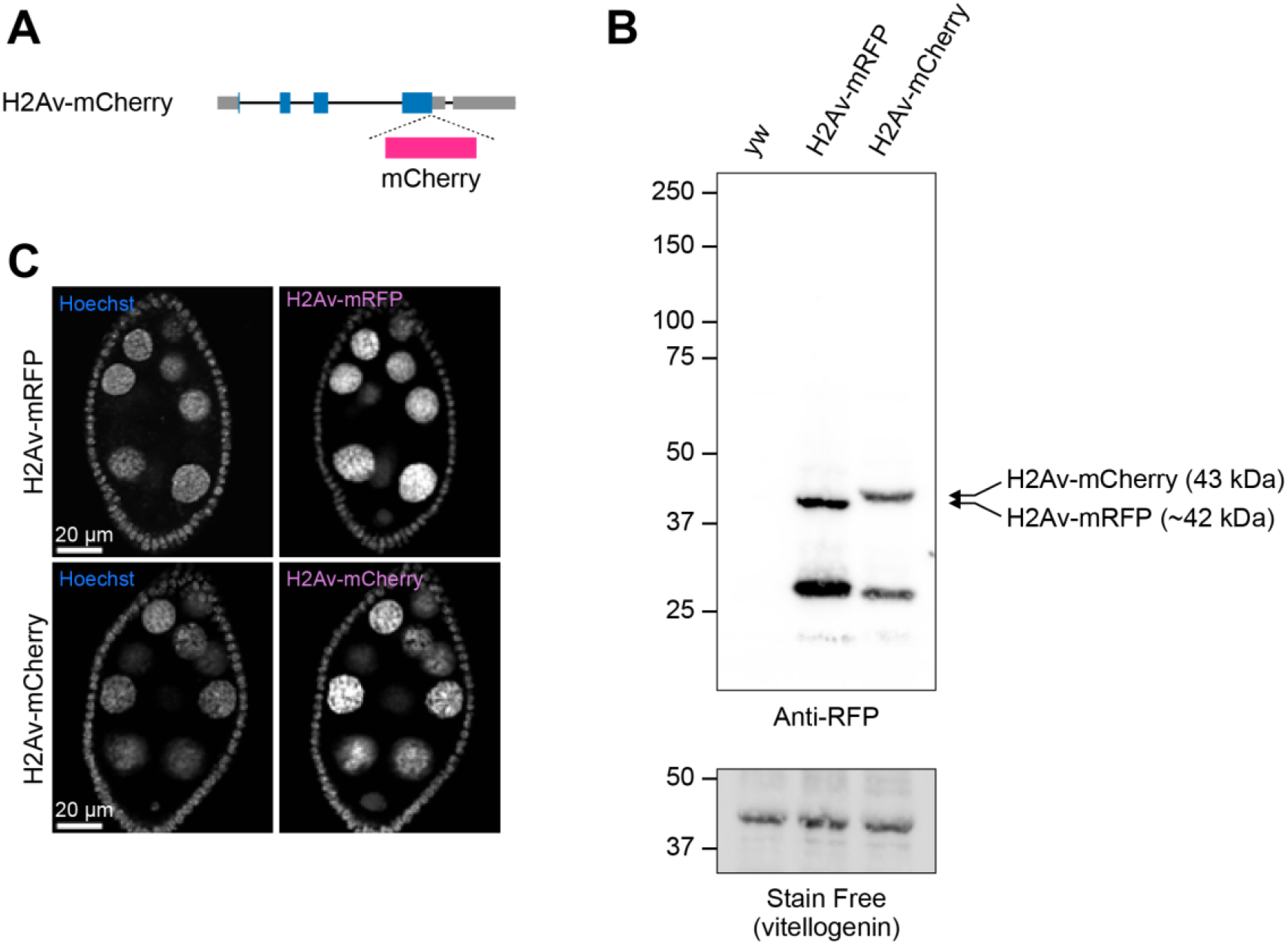
Generation of an endogenously-tagged H2Av-mCherry line. (A) Schematic of the endogenous H2Av locus tagged with mCherry using CRISPR-Cas9 mediated genome editing. (B) Western blot of the endogenously tagged H2Av-mCherry and a commonly used transgenic nuclear marker, H2Av-mRFP (His-RFP) (Schuh et al., 2007). We note that in addition to the predicted size of full-length H2Av-mRFP and H2Av-mCherry, lower molecular weight signals were detected, possibly representing cleaved products. (C) Fluorescence images of the H2Av-mRFP and the endogenous H2Av-mCherry markers in B in stage 6 egg chambers. Both markers colocalize with DNA (Hoechst) and are indistinguishable in their localization pattern.

**Figure S3:**
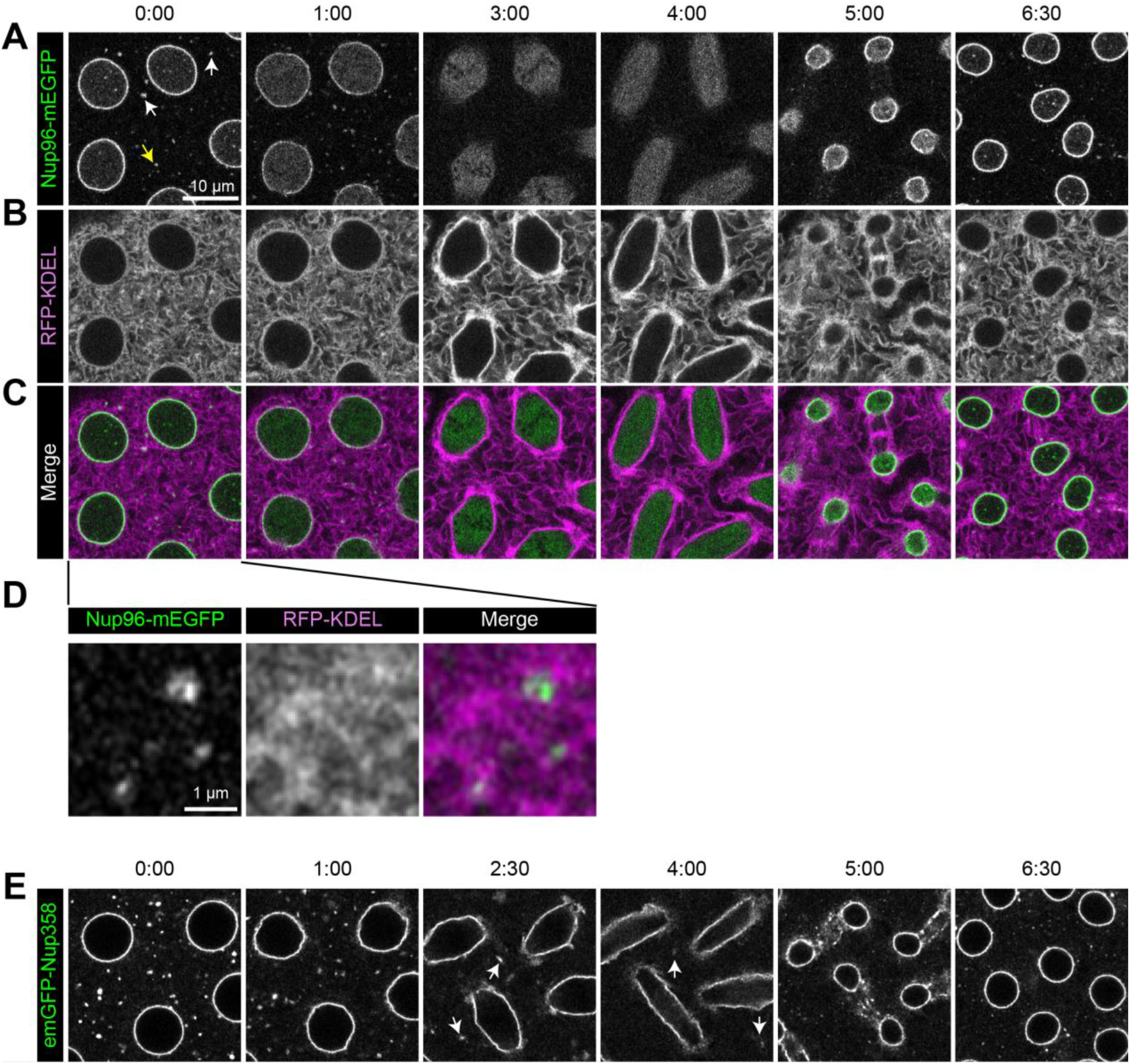
Dynamics of Nup puncta during the mitotic cycle in the early *Drosophila* embryo. (A–C) Time-lapse confocal super-resolution images of Nup96-mEGFP and matα4-tubGal4>UASp-RFP-KDEL (ER marker) in cycle 11. Arrowheads indicate cytoplasmic puncta of Nup96-mEGFP that overlap RFP-KDEL images. The Nup96-mEGFP puncta persist in interphase and prophase, dissolve in metaphase and anaphase, and reassemble in late telophase or early interphase. (D) Enlarged images of panels A–C (yellow arrowhead), showing colocalization of Nup96-mEGFP puncta and ER. (E) Time-lapse confocal super-resolution images of emGFP-Nup358. In contrast to Nup96 puncta, Nup358 puncta persist even during metaphase and anaphase (arrowheads).

**Figure S4:**
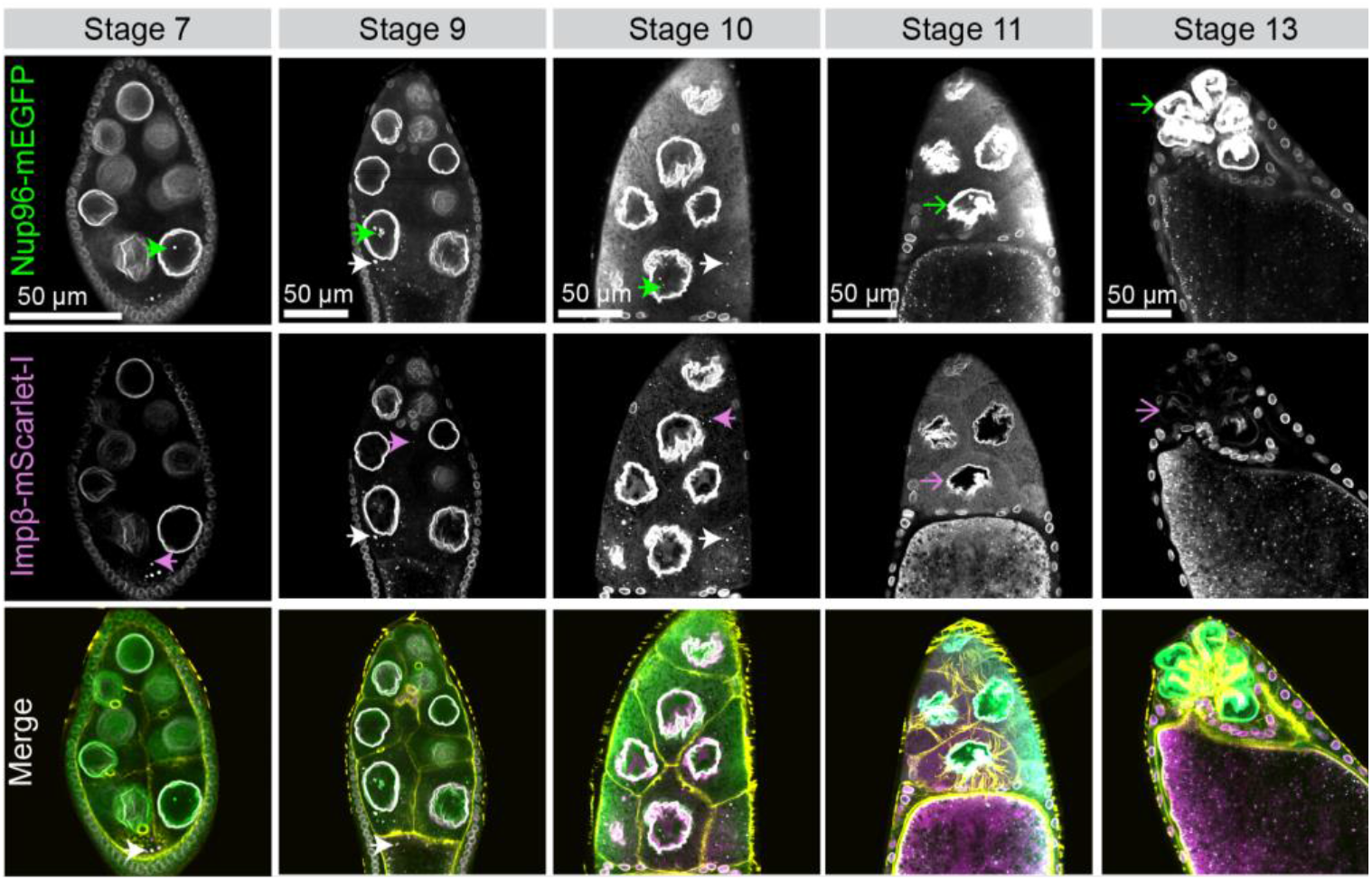
Spatio-temporal dynamics of Nup96-mEGFP and Impβ-mScarlet-I distribution during oogenesis. Nup96-mEGFP and Impβ-mScarlet-I punta appear in the oocyte cytoplasm as early as stage 5 (**Fig. 2F**). Green arrows indicate Nup96-mEGFP only puncta, magenta arrows indicate Impβ-mScarlet-I only puncta, and white arrows indicate punta with both the markers colocalized in the cytoplasm. Impβ-mScarlet-I puncta are observed in most nurse cells regardless of their position within the egg chamber, whereas Nup96-mEGFP is restricted to the nurse cells in the vicinity of the oocyte. We also observed puncta in which Nup96-mEGFP and Impβ-mScarlet-I are colocalized in the oocyte cytoplasm around stage 5 (Fig. 2F). Puncta colocalizing for both the markers in the oocyte are shown in the merged channels with phalloidin in yellow. Following nurse cell dumping (Stage 11), the Impβ-mScarlet-I levels at the nurse cell NE decrease while Nup96-mEGFP remains localized to the nurse cell NE (open arrows).

